# Actin isovariant ACT2-mediated cellular auxin homeostasis regulates lateral root organogenesis in *Arabidopsis thaliana*

**DOI:** 10.1101/2024.06.20.599894

**Authors:** Aya Hanzawa, Arifa Ahamed Rahman, Abidur Rahman

**Affiliations:** The United Graduate School of Agricultural Sciences, Iwate University, Morioka, Japan; Dept. of Plant Biosciences, Faculty of Agriculture, Iwate University, Morioka, Japan

**Author notes:** Corresponding authors’.

## Abstract

Lateral root (LR) organogenesis is regulated by cellular flux of auxin within pericycle cells, which depends on the membrane distribution and polar localization of auxin carrier proteins. The correct distribution of auxin carrier proteins relies on the intracellular trafficking of these proteins aided by filamentous actin as a track. However, the precise role of actin in lateral root development is still elusive. Here, using vegetative class actin isovariant mutants, we revealed that loss of actin isovariant ACT8 led to an increase in lateral root formation. The distribution of auxin within pericycle cells was altered in *act8* mutant, primarily due to the altered distribution of AUX1 and PIN7. Interestingly, incorporation of *act2* mutant in *act8* background (*act2act8*) effectively nullified the LR phenotype observed in *act8* mutant, indicating that ACT2 plays an important role in LR development. To explore further, we investigated the possibility that the *act8* mutant’s LR phenotype and cellular auxin distribution resulted from ACT2 overexpression. Consistent with the idea, enhanced lateral root formation, altered AUX1, PIN7 expression and auxin distribution in pericycle cells were observed in ACT2 overexpression lines. Collectively, these results suggest that actin isovariant ACT2 but not ACT8 plays a pivotal role in regulating source to sink auxin distribution during lateral root organogenesis.

## Introduction

Auxin, a major plant hormone, essentially regulates all aspects of plant growth and development (Davies, 1995). Proper function of auxin depends on its biosynthesis and intracellular distribution through auxin transport proteins that facilitates the formation of optimal auxin gradient required for cell division and various organ development (Malamy, 2005; Fukaki and Tasaka, 2009; Michniewicz et al., 2007; Vanneste and Friml, 2009). Lateral root (LR) organogenesis is one of the auxin-mediated developmental processes regulated by local auxin accumulation in pericycle cells (Laskowski et al., 2008; Péret et al., 2009; Lewis et al., 2011). Every single pericycle cell has the ability to be a founder cell depending on the available auxin concentration in that pericycle cells (De Smet et al., 2007; Dubrovsky et al., 2008). Lateral root formation consists of several developmental stages. In stage I, anticlinal and asymmetrical division of either single or pairs of pericycle founder cells create the single layer of LR primordia. Then, the cells divide periclinally to make an inner and outer layer of primordia (Stage Ⅱ). In stage Ⅲ to Ⅶ, both anticlinal and periclinal divisions happen together, and form a dome-shaped primordium. Finally, it emerges from the parental root (Stage Ⅷ). From stage Ⅷ, the lateral root starts to elongate and forms a primary root like structure (Malamay and Benfey, 1997).The developmental process from the first anticlinal cell division to the formation of a mature lateral root largely depends on the distribution of auxin in specific tissues (Malamy and Benfey, 1997; De Smet et al., 2007; Parizot et al., 2008; Perét et al., 2009; Stoeckle et al., 2018).

Optimal auxin gradient in pericycle cells required for LR initiation and development depends on the intracellular auxin transport modulated by the proper membrane localization of auxin transport proteins (Muday and DeLong, 2001). In roots, two active polar auxin transport streams exist, rootward transport from an auxin source in the shoot to basal sink tissues such as the root, and shootward transport, from the root tip toward the root-shoot junction (Baskin et al., 2010). This polar auxin transport is primarily regulated by specific set of transport proteins, auxin influx carriers, AUX (AUXIN RESISTANT1)/LAX (LIKE AUX1) protein family, and efflux carriers, PIN-FORMED protein family (Vanneste and Friml, 2009; Baskin et al., 2010; Brumos et al. 2018). AUX1 is a well-known auxin influx carrier located in the lateral root cap cells (LRC), epidermal cell in the root meristem and phloem cells of root vascular tissues (Marchant et al., 2002; Swarup et al., 2004; Swarup et al., 2008). The *aux1* mutant displays reduced auxin levels in LRC cells, which inhibits pre-branch sites and LR formation, suggesting that AUX1 plays an important role in LR positioning, initiation, and emergence (Casimiro et al., 2001; Marchant et al., 2002; De Smet et al., 2007; Xuan et al., 2016). The membrane-localized auxin efflux carrier PIN-mediated local auxin gradient is also essential for LR primordia formation (Benková et al., 2003), which is substantiated by the observation that mutations in multiple *pins* (*pin1 pin3*, *pin1 pin3 pin7*, *pin1 pin4 pin7*, or *pin1 pin3 pin4 pin7*) resulted in altered LR primordia development (Blilou et al. 2005; Laskowski, 2008; Lewis et al., 2011).

Proper cellular auxin gradient formation depends on the correct localization of the auxin transporter proteins to the membrane aided by the intracellular trafficking of these proteins (Muday and DeLong, 2001; Muday and Murphy, 2002). Actin filament is one of the essential factors for regulating cellular protein trafficking and cycling (Staiger and Schliwa, 1987; An et al., 1996; Shimmen and Yokota, 2004; Smith and Oppenheimer, 2005; Staiger and Blanchoin, 2006; Kandasamy et al., 2009; Pollard and Cooper, 2009). Not surprisingly, intracellular trafficking of both PIN1 and PIN2 had been shown to be actin dependent (Geldner et al., 2001, Rahman et al., 2007), highlighting the role of actin in optimal auxin gradient formation.

*Arabidopsis thaliana* has eight subclasses in its actin gene family which can be divided into vegetative class and reproductive class according to their ancient gene families and functions (Mclean et al., 1990; Meager et al., 1999; Kandasamy et al., 2009). Three vegetative class actin isovariants, ACT2, ACT8 and ACT7, are very similar and their homology of amino acid sequence is over 90% (Supplementary Figure 1). Although vegetative actin isovariants are highly homologous, their loss-of-function mutants represent unique phenotypes. For instance, *act2-1* mutant shows severely suppressed root hair growth (Kandasamy et al., 2009), consistent with the report that ACT2 is essential for root hair tip growth (Gilliland et al., 2002 and Ringli et al., 2002). Primary root length growth has been shown to be affected only in *act7-4* mutant but not in *act2* or *act8* mutant (Numata et al., 2022). Loss of ACT7 causes the depolymerization of actin in root that results in small meristem and short primary root (Gilliland et al., 2002; Kandasamy et al., 2009; Numata et al., 2022), indicating that ACT7 plays an important role in determining primary root development and meristem size. These findings suggest that each actin isovariant has a specific function to modulate cellular and developmental processes.

The initial asymmetric cell division required for lateral root primordia initiation is primarily dependent on the formation of proper auxin gradient in pericycle cells mediated by correct localization of auxin transporter proteins that is effectively linked with actin mediated protein trafficking. However, the role of actin in LR development remains elusive. Using vegetative class actin isovariant mutants, we tried to understand the role of actin isovariants in LR developmental process as well as intracellular auxin distribution. Our results revealed that actin isovariant ACT2 plays a pivotal role in regulating source to sink auxin distribution during lateral root organogenesis. We also demonstrated that ACT2 regulates the auxin distribution in the pericycle cells by modulating the expression of auxin influx carrier AUX1 and efflux carrier PIN7.

## Materials and Methods

### Plant materials

All lines used in this study are in the Columbia (Col-0) background of *Arabidopsis thaliana*. *act2- 1*, *act8-2*, and *act7-4* and double mutant, *act2act8* were gifts from R. Meagher (University of Georgia, Athens, Georgia, Kandasamy et al., 2009). The transgenic line harboring *DR5:GUS* (Ulmasov et al., 1997) was obtained from Jane Murfett and Tom Guilfoyle (University of Missouri, Columbia, MO, USA). Col-0 and PIN1-GFP were obtained from the Arabidopsis Biological Resource Center (Columbus, OH, USA). PIN2-GFP (Xu and Scheres, 2005) and YFP-AUX1(Swarup et al., 2007) were gifts from B. Scheres (Wageningen University, The Netherlands), and Malcolm Bennett, University of Nottingham, UK). PIN7-GFP (Vieten et al., 2005; Laskowski et al., 2008) was obtained from Gloria Muday (Wake Forest University, NC, USA, Transgenic lines of *DR5-GUS*, PIN1-GFP, PIN2-GFP, PIN7-GFP, YFP-AUX1 in actin mutants background were generated by crossing, and F3 homozygous lines were used for the experiments. Two-three independent homozygous lines for the mutation and expressing the GFP or GUS reporter were identified by screening for fluorescence, GUS assay, and seedling phenotype.

### Growth conditions

Seeds were sterilized in 20% kitchen bleach (Haita, Coop Clean Co., Japan) for 10 min and washed three times in sterilized distilled water. Surface sterilized seeds were placed in round, 9 cm Petri plates on modified Hoagland medium (Baskin and Wilson, 1997) containing 1 % w/v sucrose, 1 % w/v agar (Difco Bacto agar, BD laboratories, USA). Two days after stratification at 4°C in the dark, plates were transferred to a growth chamber (LH-240S, NK System, Japan) at 23°C under continuous white light at an irradiance of 80–90 mmol m^-2^ s^-1^. The seedlings were grown vertically for 5 or 7 days.

### Root growth assay

5-day-old seedlings were transferred to new Hoagland plates and kept at 23°C (LH-240S, NK System, Japan) for 48 h under continuous white light at an irradiance of 80-90 mmol m^-2^ s^-1^. All phenotypic images were taken by Canon EOS Kiss X10 (Canon, Japan) without flash and using micro- focus function. Root elongation was measured using ImageJ software (https://imagej.net/ij/). The number of lateral roots was quantified with the stereoscopic microscope, Nikon, Diaphot TMD equipped with a digital camera control unit, Nikon DS-L2 (Nikon, Japan).

### Live cell imaging

To image GFP and YFP, 5-day-old seedlings were incubated at 23℃ under continuous light for two days on modified Hoagland medium as described earlier. After mounting on a large cover glass (55 mm), the roots were imaged with x20 or x40 objectives using a Nikon laser-scanning microscope (Eclipse Ti equipped with NikonC2 Si laser-scanning unit, http://www.nikon.com/). The representative images and quantification were obtained from at least three biological replicates. Images were acquired using the same confocal settings for each group of experiments. Fluorescence areas were measured by drawing a region of interest in the images obtained from live-cell imaging using Image J software.

### GUS staining

GUS staining was performed as described earlier (Okamoto et al., 2008). In brief, 7-day-old vertically grown seedlings were used for GUS assay. Seedlings were transferred to GUS staining buffer (100 mM sodium phosphate pH 7.0, 10 mM EDTA, 0.5 mM potassium ferricyanide, 0.5 mM potassium ferrocyanide, and 0.1 % Triton X-100) containing 1 mM X-gluc, and incubated at 37 °C in the dark for 24h. For cell clearing, Visikol (Visikol, Inc., USA) was used as per manufacturer’s instruction. The roots were imaged with a light microscope (Nikon Diaphot) equipped with a digital camera control unit (Digital sight, DS-L2; Nikon).

### Gene expression analysis

5-day-old vertically grown *Arabidopsis* seedlings were transferred to agar plates and incubated at 23℃ for 48 hours under light condition. RNA was extracted from the root tissue using Plant RNA extraction kit (Qiagen, USA) and tested for quality and quantity. Each RNA concentration was normalized with RNase free water, and 500 ng RNA was applied to synthesize single standard cDNA using Rever Tra Ace qPCR RT master mix (Toyobo, Japan).

Quantitative PCR reactions were performed using the Takara TP-850 thermal cycler (Takara Bio, Japan) and THUNDERBIRD® SYBR™ qPCR Mix (Toyobo, Japan). Data were obtained from three biological replicates.

Since *ACT2* and *ACT8* ORFs are highly homologous, to distinguish the expression of *ACT2* from that of *ACT8*, the specific primer for *ACT2* was designed using a unique sequence region of its 3’ UTR. Similarly, *ACT8* specific primer was also designed in its 5’UTR which is different from the sequence of *ACT2*.

Primer sequences used to analyze *ACT2* expression are as follows:

Forward primer GCCAACAGAGAGAAGATGAC Reverse primer AGACACACCATCACCAGAAT

Primer sequences used to analyze *ACT8* expression are as follows:

Forward primer CACTCGGTTTATTTCTTCTCCCC Reverse primer ACAAGAACCACGACGATGAAG

### Construction of Binary Vectors and Plant Transformation

To clone the *ACT2* and *ACT8* cDNA, the primers were designed using Oligo Analyzer and Bio Labs Tm Calculator (https://tmcalculator.neb.com/#!/main).

Primers used for *ACT2* cloning are as follows:

Forward primer CACCATGGCTGAGGCTGATGATATTC Reverse primer GAAACATTTTCTGTGAACGATTCCTG Primers used for *ACT8* cloning are as follows:

Forward primer CACCATGGCCGATGCTGATGACATTC Reverse primer GAAGCATTTTCTGTGGACAATGCCTG

The *ACT2* and *ACT8* cDNA were PCR amplified and transformed into pENT vector using TOPO cloning according to manufacturer’s protocol (Life Technologies, USA). The *ACT2* gene in the construct was confirmed by DNA sequencing. The resulting constructs were transformed into *Agrobacterium tumefaciens* GV3101 for plant transformation. Col-0 plants were transformed by the floral dip method (Clough and Bent, 1998). The T1 seedlings were germinated on agar plates containing Hoagland, screened with 50µg/ml kanamycin to select transgenic plants and 100µg/ml Cefotaxime to inhibit the growth of agrobacterium, and examined for alterations in morphology. The expression level of each ACT2 and ACT8 overexpression lines were quantified by quantitative PCR.

### Western blotting

Total protein extracts were obtained from the 7-day-old whole seedlings. The whole seedlings were homogenized in sodium dodecyl sulfate-polyacrylamide gel electrophoresis (SDS–PAGE) sample buffer. A total of 50 mg protein was resolved by SDS–PAGE, transferred to Immobilon-P PVDF membrane, and probed with [mAb13a] mouse antibody, an actin subclass I specific antibody (1:2000, Kerafast, USA). Goat anti-mouse HRP antibody was used as a secondary antibody (1:5000, Jackson Immuno Research, USA). Proteins were detected with a luminoimage analyzer (AE-6972C; ATTO, https://www.atto.co.jp) using Amersham ECL prime western blotting detection reagent kit (GE Healthcare, https://www.gehealthcare.com), as described previously (Rahman et al., 2010). Band intensity was quantified using ImageJ (https://imagej.nih.gov/ij/) software. Data were obtained from three biological replicates.

### Sequence Comparison and Phylogram

Comparison of amino acid sequences of ACT2 splicing variants 1, 2, 3, and ACT8 were performed using Clustal Omega (https://www.ebi.ac.uk/Tools/msa/clustalo/). For constructing phylogenetic tree, sequences of actin isovariants were aligned in Clustal W (https://www.genome.jp/tools-bin/clustalw) using default settings.

### Statistical analysis

Results are expressed as the means ± SE. from the appropriate number of experiments as described in the figure legends. A two-tailed Students *t*-test was used to analyze the statistical significance.

## Results

### Loss of ACT8 enhances lateral root formation

Since the vegetative actin class plays important roles during early developmental stage of plants (Numata et al., 2022), in the current study, we focused only on the vegetative actins, and performed root phenotypic analyses of actin single mutants, *act2-1, act8-2, act7-4,* and double mutant *act2act8* (Figure 1 A). Consistent with previously published results, we found that loss of ACT2 and loss of ACT8 did not alter the primary root elongation, while loss of ACT7 severely inhibited the primary root elongation (Figure 1, Numata et al., 2022). Surprisingly, *act8-2* mutant showed a clear visible LR phenotype (Figure 1 A), where more LR were apparent compared with the wild-type. Quantitative analyses expressing the total LR number and LR density (LR number/ cm root length) revealed that *act8*-2 produces approximately two-fold more LR compared with wild-type (Figures 1 C and D). Although *act7-4* showed a decreased number of LRs, the density of LR did not change indicating that the observed reduction in the total LR number is due to reduced root length of *act7* (Figures 1 C and D). Collectively, these results suggest that loss of ACT8 results in promoting the LR development, and the other actin isovariants do not have any significant effect on LR production.

**Figure 1.**
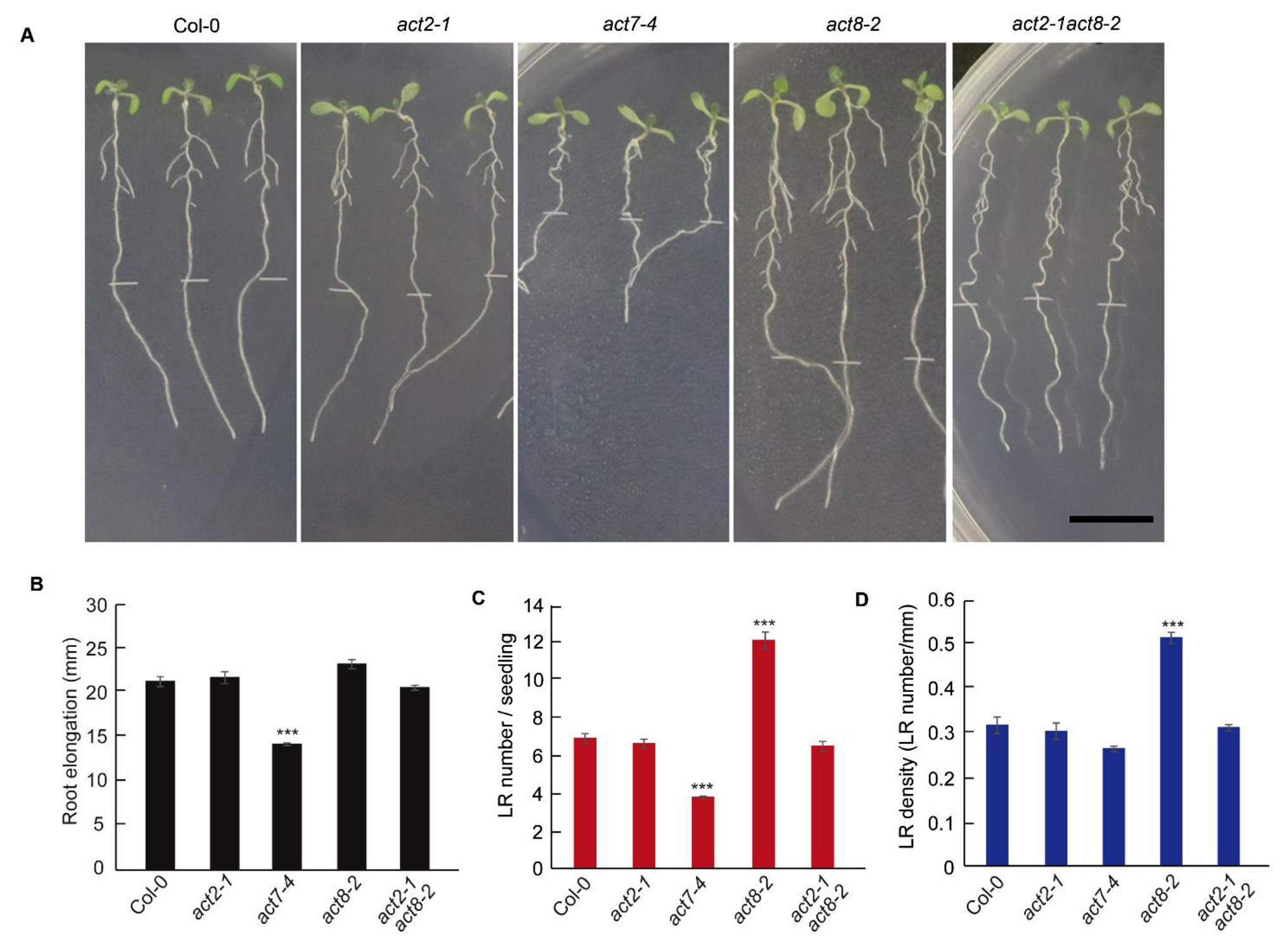
Lateral root phenotype in vegetative actin mutants A) Representative images of root phenotype of Col-0 and actin mutants. B) Primary root elongation of wild-type and actin mutants after transfer to new plates. C) Comparison of LR number in wild-type and actin mutants. D) Comparison of LR density in wild-type and actin mutants. Latreal root density represents numbers of lateral root / mm root length, which was obtained by dividing the total number of lateral roots against the whole root length. Five-day-old light-grown wild-type (Col-0) seedlings and vegetative actin isovariant mutants, *act2-1*, *act7-4*, *act8-2*, *act2act8* were transferred to new agar plates and incubated at 23°C for 2 days under continuous light. The number of lateral roots was counted under a microscope. Vertical bars represent mean ± S.E. of the experimental means from at least three to five independent experiments (*n* = 3-5), where experimental means were obtained from 6-8 seedlings per experiment. Asterisks represent the statistical significance between the means for wild-type and other genotypes as judged by Student’s *t*-test (*** P < 0.001). Scale Bar=10mm.

### Auxin homeostasis in pericycle cell is altered in *act8* mutant

For the initiation of LR, local accumulation of auxin within the pericycle cell is a prerequisite as it induces cell division (Malamy and Benfey, 1997). To visualize the auxin gradient during LR initiation in pericycle cell, auxin-sensitive marker line *DR5:GUS* was used (Ulmasov et al., 1997). *DR5:GUS* was crossed in *act8-2* mutant background, and homozygous lines were selected using both lateral root phenotype and *GUS* signal. *GUS* expression analysis revealed a clear increase in the number of auxin accumulated sites in pericycle cells in *act8-2* mutant compared with wild-type (Figure 2), confirming that the observed increase in the total LR number in *act8-2* mutant is due to enhanced LR primordia formation.

**Figure 2.**
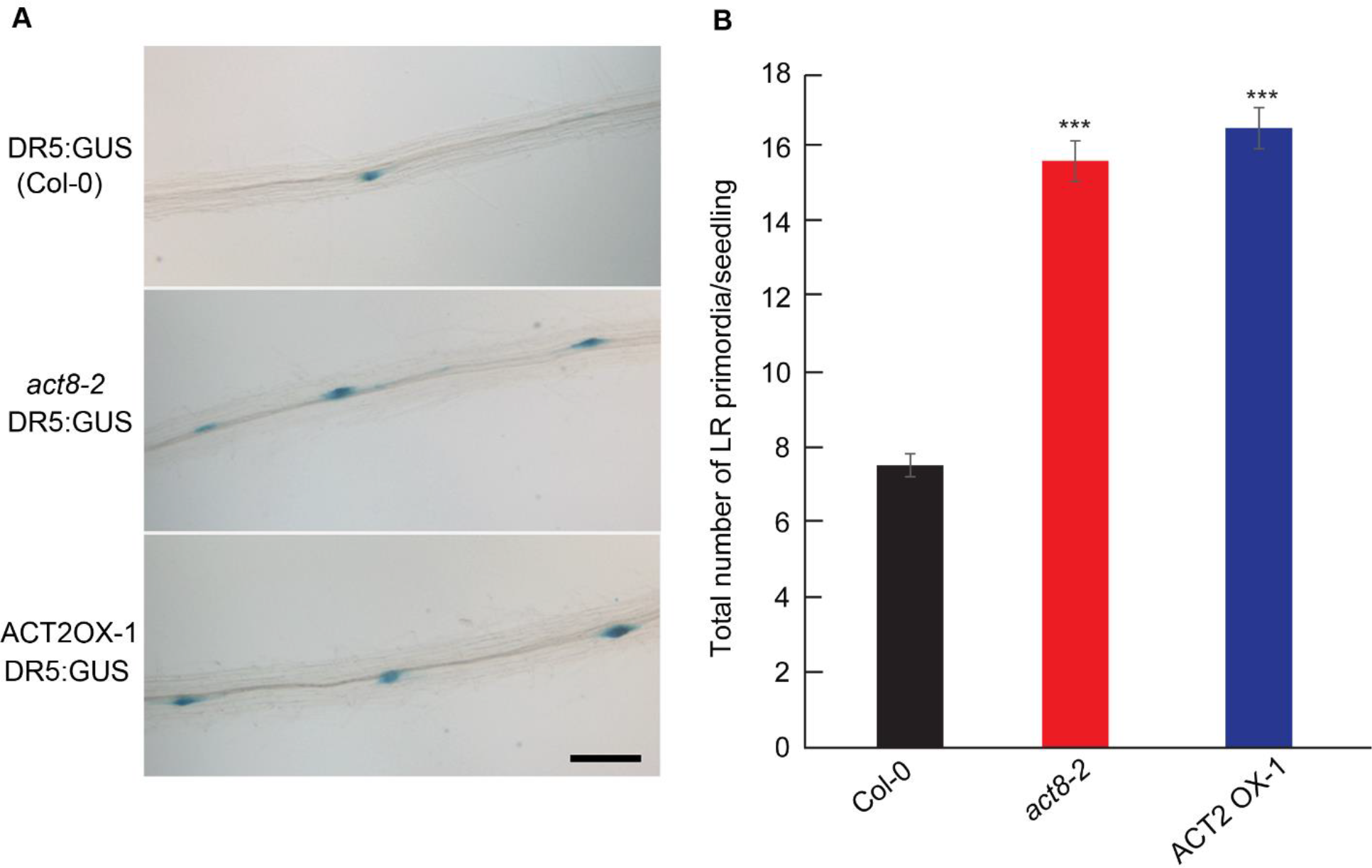
Alteration of auxin gradient in pericycle cell layer resulted in more LR initiation in *act8* mutant and ACT2 OX lines A) *DR5: GUS* signal in LR primordia located in elongation zone of Col-0, *act8-2* and ACT2 OX lines. Scale bar=100 μm. B) Quantitative analysis of the total number of LR primordia. 7-day-old *DR5:GUS* transgenic seedlings were stained in buffer containing 1mM X-gluc overnight at 37 ℃, and cleared for photography. Images are representative of 24 seedlings, stained in three separate experiments. Vertical bars represent mean ± S.E. of the experimental means from three independent experiments (*n* = 3), where experimental means were obtained from 8 seedlings per experiment. Asterisks represent the statistical significance between the means for wild-type and other genotypes as judged by Student’s *t*-test (*** P < 0.001).

### Altered auxin gradient formation in *act8* is due to altered expression of AUX1 and PIN7

The auxin response marker analysis in *act8-2* mutant revealed an increased accumulation of auxin in the lateral root founder cells (Figure 2), suggesting that loss of ACT8 may result in altered auxin transport. Both auxin uptake and efflux carriers function in coordination to ensure the optimal source to sink distribution of auxin in various cells to facilitate optimal organogenesis and growth (Marchant et al., 2002; Lewis et al., 2011)

Hence, we investigated the expression and localization of various auxin transporters, namely AUX1, PIN1, PIN2, and PIN7, that had been demonstrated to regulate auxin gradient formation during LR organogenesis (Marchant et al., 2002; Lewis et al., 2011; Omelyanchuk et al., 2016; Zhao et al., 2023). GFP or YFP tagged lines of these proteins were crossed with *act8-2*, and the homozygous plants were selected for LR phenotype and fluorescence signal.

AUX1 expression was found to be upregulated in *act8-2* mutant compared with wild-type. At the root meristem, AUX1 expression area was extended in the *act8-2* mutant, which was confirmed by quantitative analysis. Compared with wild-type, AUX1 expression was extended to additional 100 μm in *act8* mutant (Figure 3 A, B). Consistently, more AUX1 expression was observed in the pericycle cells of *act8-2* compared with wild-type (Figure 3 C). These results indicate that *act8-2* accumulates more auxin in pericycle cells through modulating its influx capacity.

**Figure 3.**
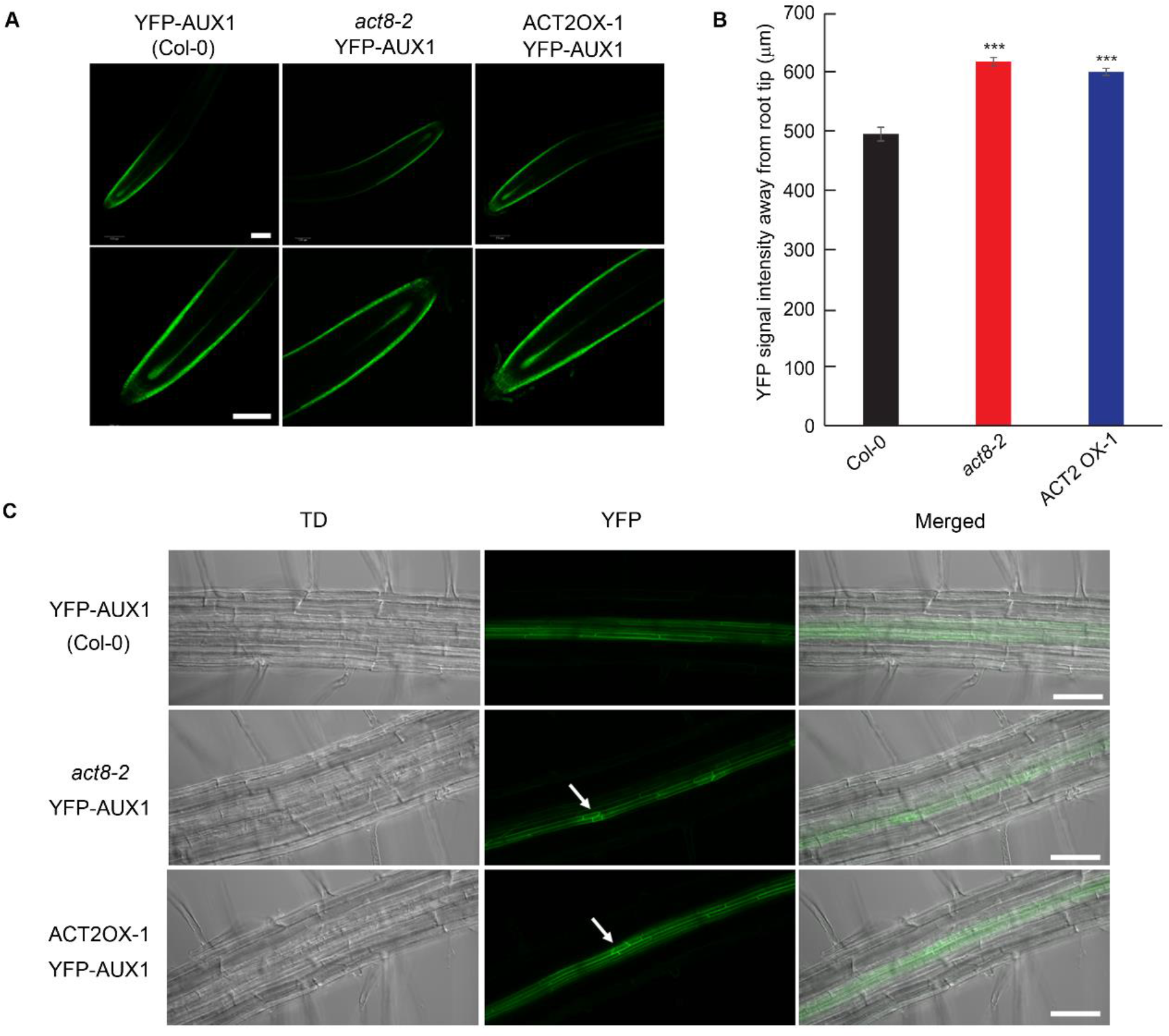
AUX1 expression is altered in *act8-2* and ACT2 overexpression line A) YFP-AUX1 expression in the root meristem of Col-0, *act8-2* and ACT2OX line. B) Quantification of AUX1 expression area in peripheral cells of Col-0, *act8-2* and ACT2OX line. The area of fluorescence signal in the meristem zone was measured by drawing a ROI using Image J. 7-day-old roots were used for imaging and the images were captured with the same confocal setting in each YFP line. The images are representative of at least 15 roots observed in three independent experiments. Scale bars=50 μm. Vertical bars represent mean ± S.E. of the experimental means from three independent experiments (n = 3), where experimental means were obtained from 5 seedlings per experiment. Asterisks represent the statistical significance between the wild-type and other genotypes as judged by Student’s t-test (*** P < 0.001).

PIN1 and PIN2 expressions in the meristem and LR initiation sites in pericycle cells were similar in wild-type and *act8-2* mutant (Supplementary Figures 2 and 3). PIN2 did not express in the LR initiation site, indicating that PIN2 is not directly involved in lateral root organogenesis (Supplementary Figure 3B). In wild-type, PIN7 shows a remarkable expression pattern where it was expressed only at the initial stage of LR development and the expression of PIN7 was turned off at the later stage of LR development (Figure 4). Interestingly, a clear increase in PIN7 expression was observed in *act8-2* mutant, which continues to express even at the later stage of LR development (Figure 4). Taken together, these results suggest that loss of ACT8 specifically affects AUX1 and PIN7 activity but not the other auxin transporter proteins.

**Figure 4:**
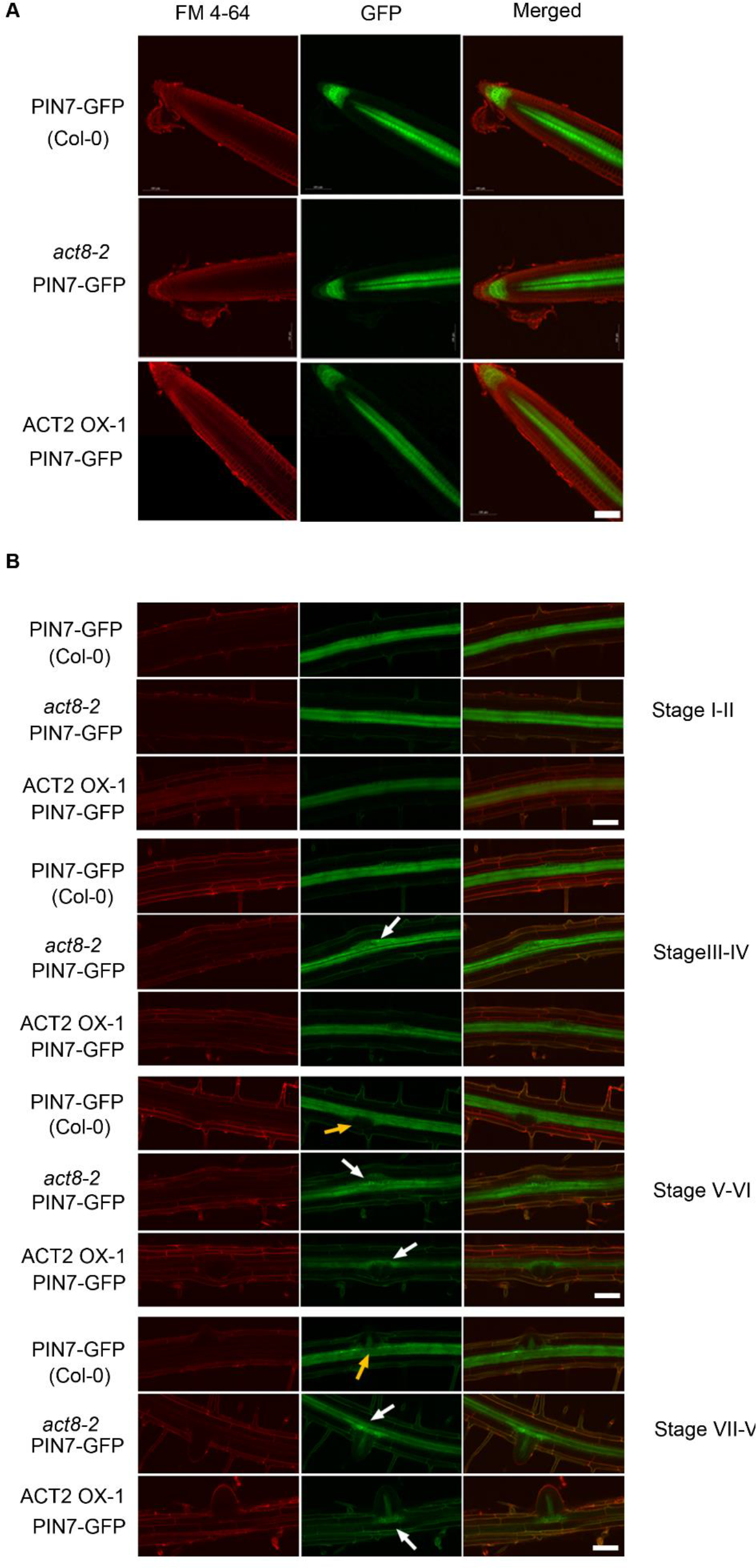
PIN7 expression is altered in the lateral root initiation sites of *act8-2* and ACT2 OX lines. A) PIN7-GFP expression in root meristem of Col-0, *act8-2* and ACT2 OX line. No significant differences were observed in the PIN7 expression between genotypes. B) PIN7-GFP expression in pericycle cells of Col-0, *act8-2* and ACT2 OX-1 line. White arrows indicate GFP fluorescent signal and yellow arrows indicate the depletion of GFP signal. The images are representative of at least 15 roots observed in three independent experiments. Scale bars=50 μm.

### Lateral root phenotype of *act8* results from overexpression of ACT2

The observation that *act8-2* mutant showed increased LR production raises a possibility that ACT8 might regulate LR development negatively in wild-type. However, this hypothesis cannot be explained as the *act8* LR phenotype is effectively nullified by the loss of ACT2 (*act2act8* double mutant showed wild-type-like LR phenotype, Figure 1). In addition, it is also difficult to characterize actin as a negative regulator for plant development since actin functions positively in various types of cellular activities including intracellular protein trafficking. Further, actin positively regulates auxin transport, which is a pre-requisite for LR development.

This raises another possibility that the observed LR phenotype in *act8* mutant may be due to the overexpression of ACT2. In fact, using the antibody that detects both ACT2 and ACT8 proteins, Kandasamy et al., (2009) demonstrated that compared with wild-type, ACT2-ACT8 protein expression was reduced to half in *act2* mutant, while expression was not changed in the *act8* mutant. This result suggests that the unaltered expression of ACT2-ACT8 protein in *act8* mutant may be due to the increased expression of ACT2.

This led us to propose the hypothesis that ACT2 may function as a positive regulator of LR induction and the observed LR phenotype in *act8* may be due to overexpression of ACT2. To test this hypothesis, first we checked *ACT2* expression in *act8-2* mutant using quantitative PCR. *ACT2* expression was found to be highly upregulated in *act8* mutant line (Figure 5 C). At the translational level, wild-type like ACT2 expression was observed in *act8* mutant line which is consistent with the results observed by (Kandasamy et al., 2009) and confirms that in *act8* mutant ACT2 is overexpressed (Figure 5 D, E). Next, we overexpressed ACT2 in Col-0 background using 35S promoter and observed whether LR development was altered in the overexpression lines. ACT2 overexpression was confirmed both at transcriptional and translational levels (Figure 5, C, D, E). At the phenotypic level, ACT2 OX lines showed more LR induction, although the primary root length was not altered (Figure 5 A, B). These results strongly support our hypothesis that ACT2 actin isovariant functions positively to regulate LR development.

**Figure 5.**
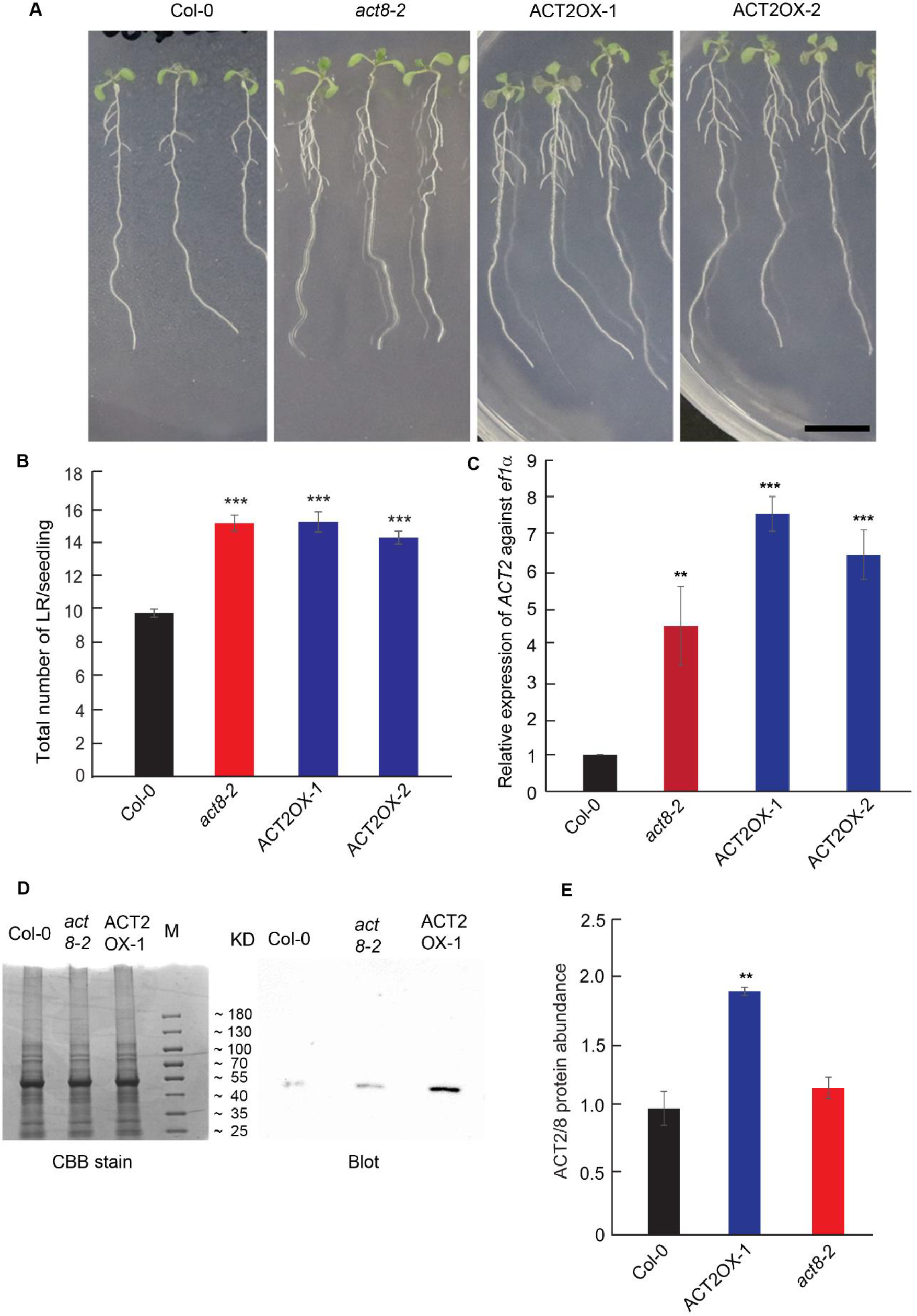
*ACT2* overexpression induces LR production A) Root phenotype of Col-0, *act8-2* and ACT2OX lines. 5-day-old seedlings were transferred into new plates containing agar Hoagland media, and incubated at 23°C for additional two days. Images are representative of 18-20 seedlings observed in three independent experiments. Bar = 10 mm. B) Number of lateral roots per seedling in each genotype. Vertical bars represent mean ± S.E. of the experimental means from three independent experiments (n = 3), where experimental means were obtained from 8 seedlings per experiment. Asterisks represent the statistical significance between Col- 0 and other genotypes as judged by Student’s t-test (**P<0.01, ***P<0.001). C) Expression of *ACT2* gene in Col-0, *act8-2* and ACT2OX lines. 7-day-old seedlings were used for RNA isolation, cDNA preparation and qRT-PCR analysis. All data were normalized against *ef1α* transcript. Vertical bars represent mean ± S.E. of the experimental means from three independent experiments (n = 3). Asterisks represent the statistical significance between Col-0 and other genotypes as judged by Student’s *t*-test: D) ACT2-ACT8 protein expression in Col-0, *act8-2* and ACT2OX-1 line. Total protein was collected from 7-day-old seedlings, resolved by the gradient (5-20%) sodium dodecyl sulfate-polyacrylamide gel electrophoresis (SDS-PAGE), and probed with monoclonal antibody against ACT2 and ACT8. Total 50 μg protein was loaded in each lane. The blot image is representative of two independent experiments. E) Relative quantification of Western blot analysis from experiment (D). Col-0 expression was considered as 1. Asterisks represent the statistical significance between the means for each genotype Student’s t-test (**P < 0.001). Vertical bars in the graph represent mean ± SE.

### ACT2 overexpression alters auxin homeostasis in the pericycle cells by altering the expression and localization of AUX1 and PIN7

In *act8-2* mutant, the altered LR phenotype was attributed to increased expression of AUX1 and PIN7. Hence, we investigated whether a similar alteration can be observed in ACT2 overexpression (ACT2OX) lines. To confirm that ACT2OX lines produce more LR primordia like *act8* mutants, auxin responsive marker *DR5::GUS* was crossed in ACT2OX lines, and homozygous lines were selected using both lateral root phenotype and GUS staining. The investigation of auxin gradient formation showed higher GUS activity in the pericycle cells in ACT2OX lines (Figure 2), confirming that ACT2 overexpression can alter auxin homeostasis in pericycle cells resulting in induction of many LRs. Next, we investigated the expression of AUX1, PIN7 as well as PIN1 and PIN2 transporters in ACT2OX lines. AUX1-YFP, PIN1-GFP, PIN2-GFP, and PIN7-GFP lines were crossed with ACT2 OX lines, and the homozygous plants were selected through screening for LR phenotype and GFP signal.

Comparison of their expression revealed that PIN1 and PIN2 expression was not altered in ACT2 OX lines, as observed in *act8-2* (Supplementary Figures 2 and 3). For AUX1 and PIN7 expression, similar expression patterns were observed in the ACT2OX lines as observed in *act8* mutant. AUX1 expression was extended in the root meristem region and enhanced in the pericycle cells (Figure 3). For PIN7 expression, no alteration was observed in the root meristem, but a clear enhancement of expression was observed in the lateral root initiation sites in pericycle cells (Figure 4). Collectively, these results strongly support the hypothesis that the observed LR phenotype in *act8* mutant is due to the overexpression of ACT2, which alters the auxin homeostasis in the pericycle cells through modulating AUX1 and PIN7 expressions and enhances the LR primordia formation.

## Discussion

LR organogenesis is a complex process that is initiated by the asymmetric anticlinal division of the pericycle founder cells followed by periclinal divisions and the whole process depends on the local accumulation of auxin and response (Malamy et al., 1997; Péret et al., 2009). Several studies revealed that the accumulation of auxin in specific founder cells is dependent on the loading and unloading of auxin regulated by both auxin influx and efflux carriers AUX1, and PIN1, PIN3, PIN4 and PIN7 (Casimiro et al., 2001; Marchant et al., 2002; Blilou et al. 2005; Laskowski, 2008; Lewis et al., 2011). Higher expression of AUX1, results in additional auxin import, creation of an auxin maximum and LR initiation (Laskowski et al., 2008). On the other hand, reduction in PIN3 and PIN7 expression at the LR initiation site strongly correlates with the efficacy of LR development (Laskowski et al., 2008; Lewis, 2011). Taken together, these results suggest that spatiotemporal expression of auxin influx and efflux carriers facilitate the LR organogenesis. Consistent with this idea, in this study we found that the observed LR phenotype in *act8* or ACT2OX lines resulted from the cellular auxin distribution in the pericycle cells regulated by the increased expression of AUX1 and PIN7 (Figures 3 and 4). In the ideal condition, the longitudinal spacing of LR plays an important role in determining the total number of LR. Earlier it was demonstrated that decrease in the expression of PIN3 and PIN7 at the LR initiation site effectively limits the region for lateral root induction (Laskowski et al., 2008). In *pin237* mutant, the longitudinal spacing of LR is diminished resulting in marked increased in lateral root density not correlated with root length. Additionally, it was also observed that when PIN3 and PIN7 expressions were not abolished, multiple primordia were formed (Laskowski et al., 2008). In both *act8-2* and ACT2OX lines, we observed an increase in both AUX1 and PIN7 expression at the lateral root initiation site of the pericycle cells and increased number of primordia (Figures 2-4). These observations are consistent with the model proposed by Laskowski et al, 2008. Increased AUX1 expression results in increased auxin influx in the pericycle founder cells and facilitates the initiation of cell division, while increased expression of PIN7 removed the regulation of longitudinal spacing of LR resulting in formation of increased number of LR. Interestingly, we did not observe any alteration in PIN1 or PIN2 expression in *act8-2* or ACT2OX lines (Supplementary Figures 2-3).

Additionally, we also observed an increase in AUX1 expression in the root meristem in both *act8- 2* and ACTOX lines (Figure 2A). Although every xylem cell has the ability to divide in response to elevated auxin level, only a limited number of pericycle cells become founder cells from where the LR initiates (Dubrovsky et al., 2001). Hence, the LRP spacing and priming model have been proposed (De Smet et al.,2007; Laskowski et al., 2008). LR priming occurs in basal meristem using the local auxin gradient aided by the shootward auxin transport system (De Smet et al., 2007; Péret et al., 2009). Further, it was demonstrated that Arabidopsis pericycle cells opposite to the xylem pole are shorter than other founder cells, suggesting that these cells undergo more divisions in the root apical meristem than other cells (Beeckman et al., 2001). We assume that increased AUX1 expression in the root meristem facilitates the priming and mitotic activity of the founder cells by increasing the local auxin gradient, and thus prepare the founder cells to be able to initiate LRP at a later stage of development. Collectively these results suggest that the altered LR phenotype observed in *act8-2* and ACTOX lines is specifically regulated by the altered expression of AUX1 and PIN7 that facilitates the priming, spacing, and ultimately the LR initiation.

Auxin gradient formation along the root depends on the expression and correct localization of the auxin transporters. The appropriate localization of the auxin transporters protein inside the cell is regulated by protein trafficking where actin plays a crucial role (Muday and Murphy 2002). Depolymerization of actin filaments disturbs the trafficking and localization of auxin influx and efflux proteins resulting in altered intracellular auxin homeostasis and development (Geldner et al., 2001, Friml et al., 2003; Kleine-Vehn et al., 2006; Rahman et al., 2007). Hence, it is logical to assume that actin may directly influence the LR developmental process along with other cellular processes. Here, we demonstrated that ACT2 plays an important role in regulating the LR developmental process. In Arabidopsis, three actin isovariants, ACT2, ACT8 and ACT7 constitute the vegetative class actin. ACT2 and ACT8 belong to the same clade and show 99.7% homology, while the homology between ACT2/ACT8 and ACT7 is around 93% (Supplementary Figure 1). Although these actin isovariants show high homology, they have been shown to regulate distinct developmental processes. For instance, ACT2 and ACT8 regulate bulge site selection, root hair growth, and leaf morphology (Ringli et al., 2002; Nishimura et al., 2003; Kandasamy et al., 2009, Vaškebová et al., 2018), while ACT7 regulates germination, root growth, meristem development, epidermal cell specification, root architecture, and root thermomorphogenesis (Gilliland et al., 2003; Kandasamy et al., 2009; Parveen and Rahman, 2021, Numata et al., 2022). Additionally, we recently demonstrated that ACT7 but not ACT8 or ACT2 plays a crucial role in maintaining the structural integrity of actin filament. Loss of ACT7 results in fragmented and depolymerized actin that affects the expression and trafficking of a subset of auxin efflux carriers mainly PIN1, PIN2, and PIN7, and subsequently the polar transport mediated auxin gradient at the root tip resulting in reduced cell division and shorter root meristem (Numata et al., 2022).

Here, we demonstrated that ACT2 plays an important role in regulating the LR organogenesis through modulating the expression of AUX1 and PIN7. The LR phenotype observed in the *act8-2* mutant was found to be due to the overexpression of ACT2. This finding is supported by several lines of evidence. The LR phenotype of *act8-2* is completely nullified in the *act2act8* double mutant, suggesting that ACT2 plays an important role in developing LR phenotype in *act8-2* mutant background (Figure 1). Overexpression of ACT2 in *act8-2* mutant was confirmed both at transcriptional and translational level (Figure 5 C-E). The combined expression of ACT2 and ACT8 was not reduced in *act8-2* mutant background although in *act2-1* mutant background the expression level was reduced to half (Figure 5 D-E, Kandasamy et al., 2009), confirming that ACT2 is overexpressed in *act8-2*. The observed LR phenotype in *act8-2* is truly due to over expression of ACT2 was confirmed by overexpressing ACT2 in wild-type background. ACT2 over expression lines showed the same LR phenotype like *act8-2* and also affected the expression of AUX1 and PIN7 in a similar fashion (Figures 2-5). The specific role of ACT2 in LR developmental regulation was further confirmed by analyzing the LR phenotype in ACT8OX lines. Overexpression of ACT8 did not alter the LR phenotype, confirming that ACT2 but not ACT8 regulates the LR organogenesis in Arabidopsis (Supplementary Figure 4).

Collectively these results strongly suggest that actin isovariant ACT2 plays a crucial role in Arabidopsis LR organogenesis through altering AUX1 and PIN7 expression and thereby modulating intracellular auxin homeostasis. However, there are still some questions that remain unanswered. First, *act2act8* double mutant did not show a reduced LR number (Figure 1). This indicates that ACT2 function is possibly compensated by other actin isovariant. The second question is why ACT2 and ACT8 show differential activities for LR organogenesis. Considering that *ACT2* and *ACT8* genes encode almost identical proteins with only one amino acid difference in the N terminal (glu3→asp3, respectively), it is difficult to explain the differential activity by protein function. Further, in contrast to ACT2, the overexpression of ACT8 did not induce LR phenotype (Figure 5, Supplementary Figure 4), suggesting there may be some other regulatory mechanisms distinguishing the function of these two isovariants. Previously it was demonstrated that 5’ flanking sequence of ACT2 and ACT8 show high level of nucleotide sequence divergence (An et al., 1996). Expression of the 5’ flanking regions of *ACT2* and *ACT8* under *GUS* reporter gene revealed a strong expression of *ACT2* in all vegetative tissue while *ACT8* expression was weak and expressed only in a subset of tissue expressing *ACT2* (An et al., 1996). Further, it was demonstrated that mutation of the element -774 in Arabidopsis *ACT2* 5’ regulatory region reduced *ACT2* expression to 60%, while mutation of the elements -566, -576, -597, -667, and -858 increased their expression 84-123% (An et al; 2010). Similar results were obtained with *Physcomitrella patens* (An et al., 2010). These results strongly suggest that the 5’ region of *ACT2* has a strong regulatory role for its expression. Perhaps, this difference in 5’ regulatory regions in *ACT2* and *ACT8* may explain the reasons for their differential activities for LR organogenesis. Answering these questions awaits further experimentation.

## Acknowledgement

The authors thank Rich Meagher (University of Georgia, Athens, USA), Tom Guilfoyle (University of Missouri, Columbia, MO, USA), B. Scheres (Wageningen University, The Netherlands), Malcolm Bennett (University of Nottingham, UK), and Gloria Muday (Wake Forest University, NC, USA for sharing the materials. We also thank Marika Yamauchi for technical assistance.

## Author contributions

AR designed the experiments; AH and AAR performed the experiments; AR supervised the experiments; all the authors analysed the data; AH, AAR and AR wrote the manuscript.

## Funding

This work was funded by JSPS Kakenhi (Grant Number 23012003 and 20H02945 to AR).

## Conflict of interest

The authors declare no conflict of interest. The funders had no role in the design of the study; in the collection, analyses, or interpretation of data; in the writing of the manuscript; or in the decision to publish the results.

## Notes

### Competing Interest Statement

The authors have declared no competing interest.

