## Supplemental figure for "Actin isovariant ACT2-mediated cellular auxin homeostasis regulates lateral root organogenesis in *Arabidopsis thaliana*"

**A**

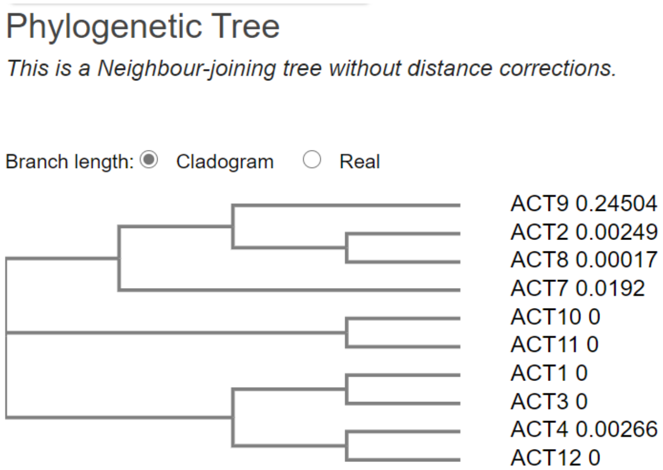


**B**

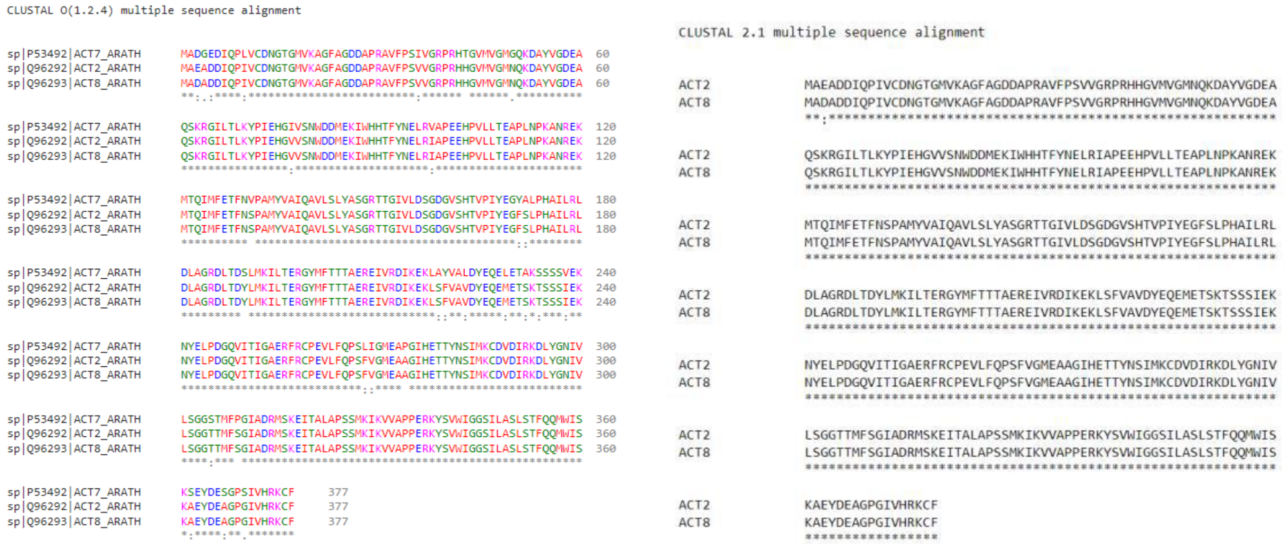


**C**

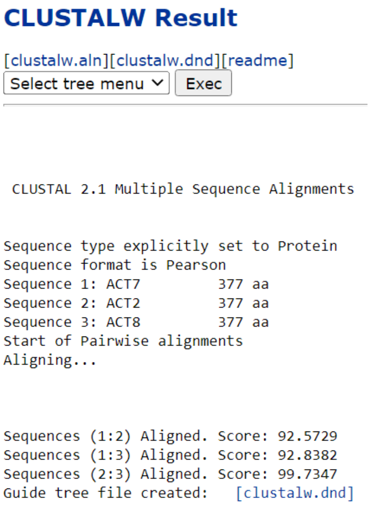


**Supplemental Figure 1:**
A) Phylogenetic analysis of Actin proteins in *Arabidopsis thaliana*.
The tree was built using Clustal Omega
B-C) Multiple Sequence alignment and homology score between ACT2, ACT8 and ACT7. Please note that the homology between ACT2 and ACT8 is 99%, between ACT2 and ACT7 is 92.5%, and between ACT8 and ACT7 is 92.8%.

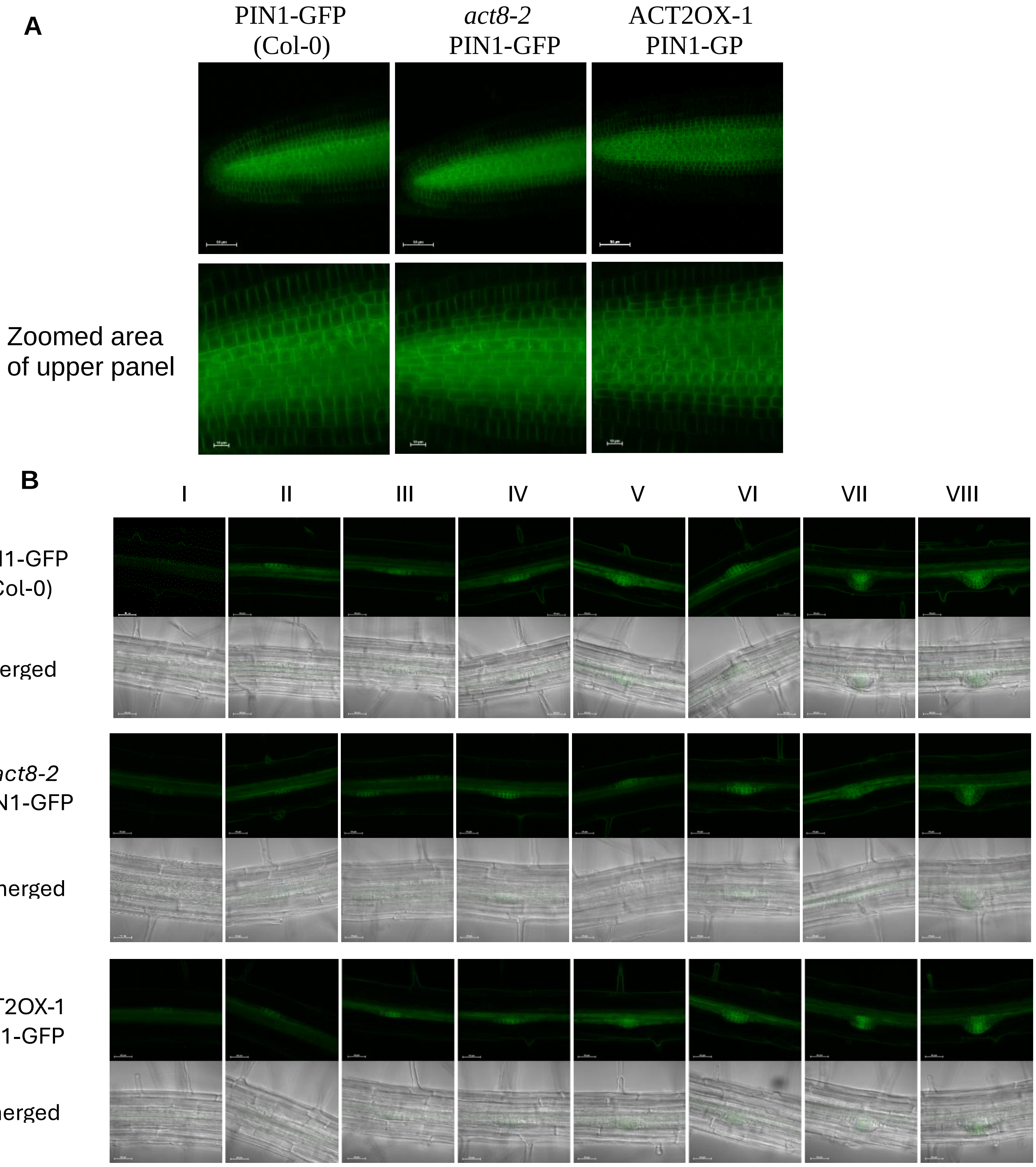


**Supplemental Figure 2:** PIN1-GFP expression in different genotypes.
A) PIN1-GFP expression in root meristem of Col-0, a*ct8-2* and ACT2 OX lines. Scale bar=50 µm. B) PIN1-GFP expression in different stages of lateral root development of Col-0, *act8-2* and ACT2 OX lines Scale bar=50 µm.

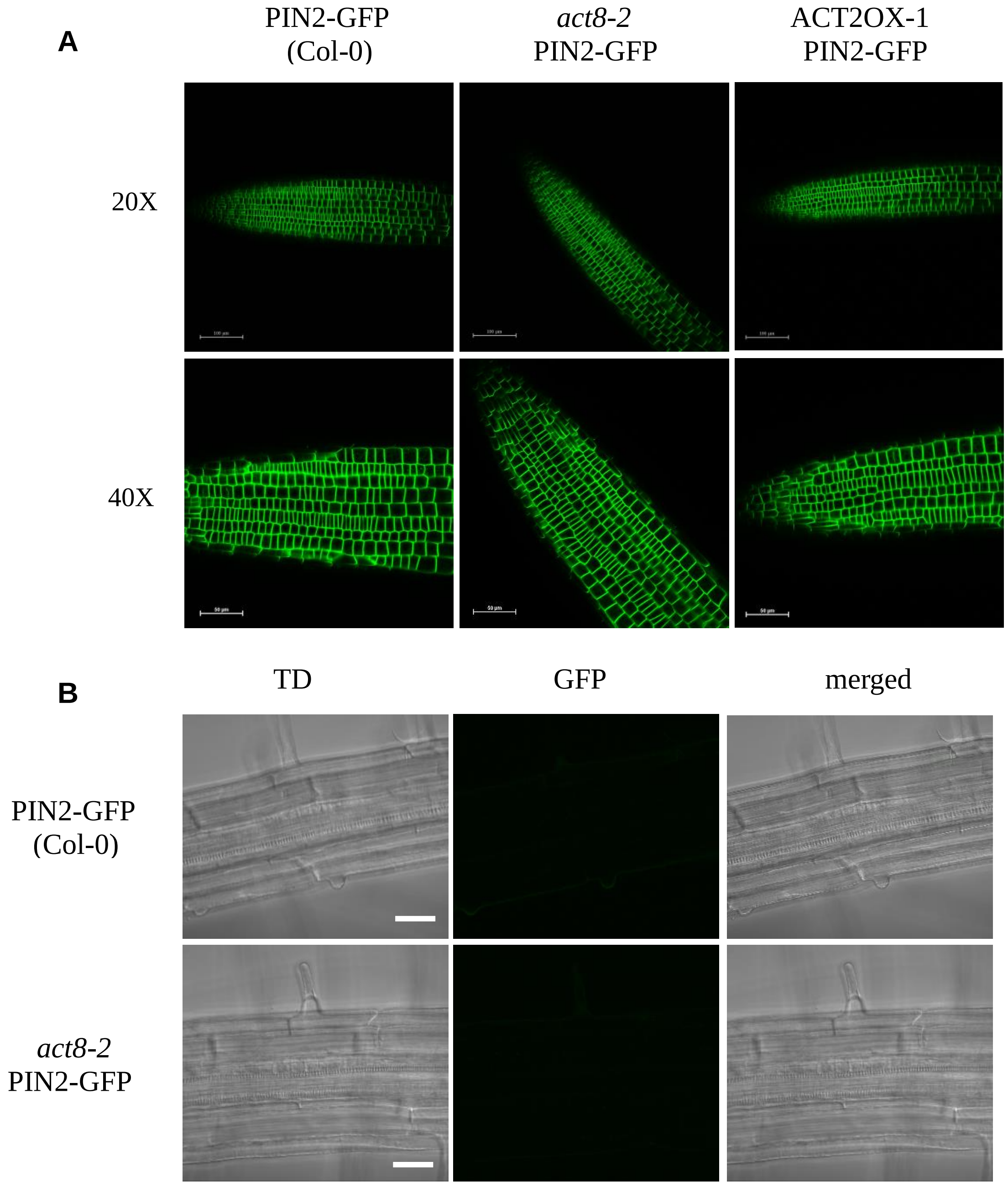


**Supplemental Figure 3:** PIN2-GFP expression in different genotypes.
A) PIN2-GFP expression in the root meristem of Col-0, *act8-2* and ACT2 OX lines. Scale bar=100 µm and 50 µm in 20x and 40x, respectively. B) PIN2-GFP expression in pericycle cells of Col-0 and *act8-2* Scale bar=50 µm. Please note that PIN2 did not express at all in pericycle cells.

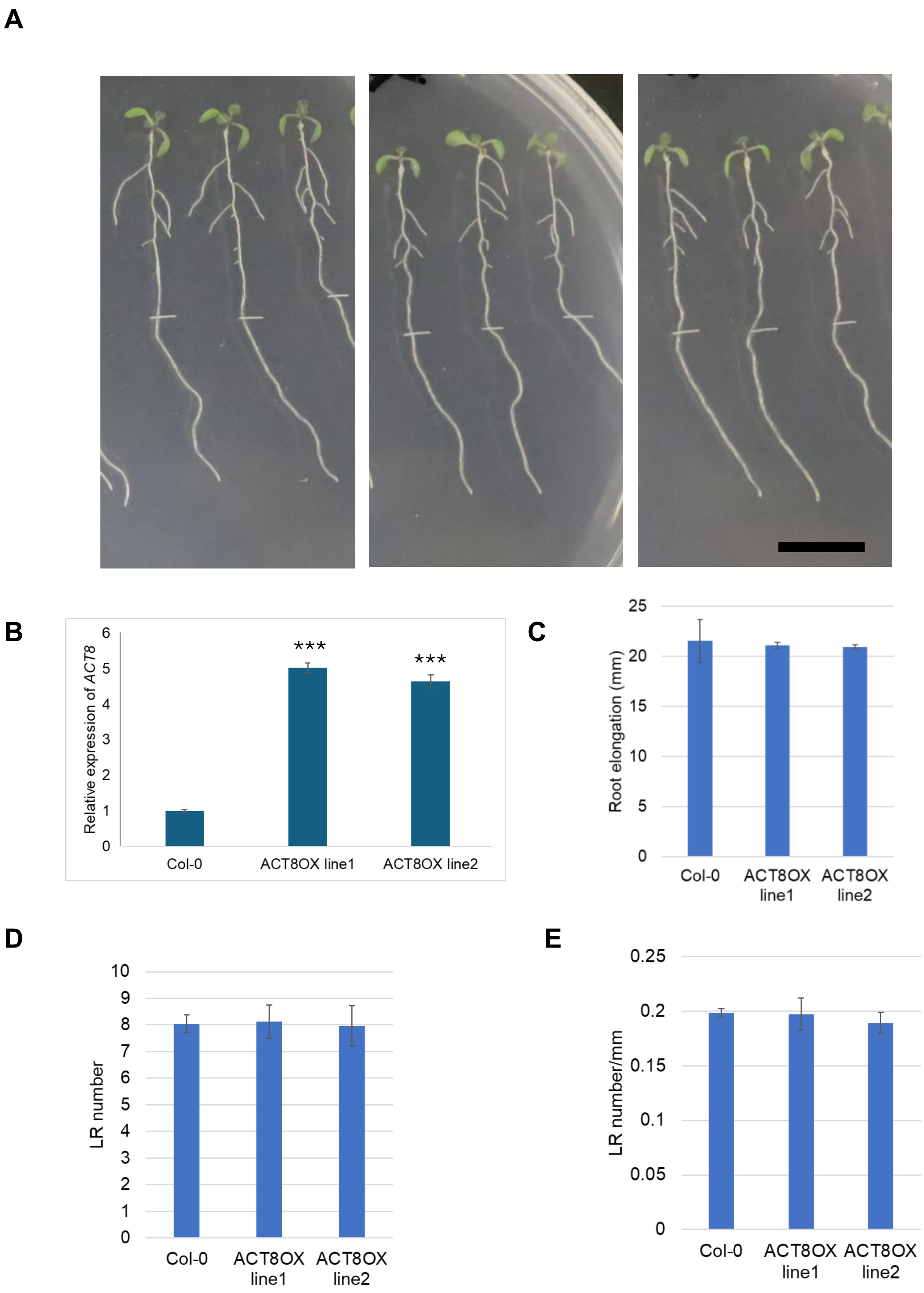

A
B
C
D
E
Relative expression of ACT8
***
***
Col-0
ACT8OX line1
ACT8OX line2
Root elongation (mm)
LR number
LR number/mm

**Supplemental figure 4:** Lateral root phenotype in ACT8 OX lines.
Five-day-old light-grown wild-type (Col-0) seedlings and ACT8 OX lines were transferred to new agar plates and incubated at 23°C for 2 days. The number of lateral roots was counted under a microscope.
A) Root phenotype of wild-type and ACT8 OX lines. Scale bar=10 mm. B) Expression of *ACT8* gene in Col-0 and ACT8OX lines. 7-day-old seedlings were used for RNA isolation, cDNA preparation and qRT-PCR analysis. All data were normalized against *ef1α* transcript. Vertical bars represent mean ± S.E. of the experimental means from three independent experiments (n = 3). Asterisks represent the statistical significance between Col-0 and other genotypes as judged by Student's *t*-test (***P < 0.0001). C) Primary root elongation of wild-type and ACT8 OX lines. D) Comparison of LR number in wild-type and ACT8 OX lines. E) Comparison of LR density (number/mm root) in wild-type and ACT8 OX lines. The images are representative of three biological replicates. Vertical bars in the graph represent mean± SE obtained from three replicates.
